# Wnt10b signaling regulates replication stress-induced chromosomal instability in human cancer

**DOI:** 10.1101/2025.04.23.650265

**Authors:** Alexander Haas, Friederike Wenz, Janine Wesslowski, Gary Davidson, Oksana Voloshanenko, Michael Boutros, Sergio Acebron, Holger Bastians

## Abstract

Wnt signaling pathways are involved in various developmental and tissue maintenance functions while deregulated Wnt signaling is closely linked to human cancer. Recent work revealed that loss of Wnt signaling impairs mitosis and causes abnormal microtubule growth at the mitotic spindle resulting in chromosome missegregation and aneuploidy, both of which are hallmarks of cancer cells exhibiting chromosomal instability (CIN). Here, we demonstrate that Wnt signaling specifically activated by Wnt10b is relevant in colorectal cancer cells to prevent abnormal microtubule dynamics and chromosome missegregation. Although mitosis is affected, Wnt10b signaling is required during the S phase of the cell cycle. In fact, Wnt10b signaling acts upon DNA replication stress in S phase, a condition typically associated with CIN in cancer, to prevent increased microtubule dynamics from S phase until mitosis, where they otherwise manifest in chromosome missegregation. Interestingly, replication stress-induced chromosomal breaks are also efficiently suppressed by Wnt10b. Thus, our results show that Wnt10b signaling regulates replication stress-induced chromosome missegregation and breakage, and hence, is a determinant for broad genome instability in cancer cells.

**Summary blurb:** We describe a novel role of Wnt10b signaling acting in response to DNA replication stress to suppress chromosomal breaks and mitotic errors in human cancer cells.

## Introduction

Wnt signaling pathways play central and diverse roles in development and tissue homeostasis (Albrecht et al, 2021, Nusse et al, 2017). The best investigated example is the Wnt-β-catenin pathway that can be activated for instance by Wnt3a ligand leading to stabilization and activation of the transcriptional regulator β-catenin. This, in turn, drives the expression of a multitude of target genes contributing to the various cellular functions of Wnt signaling (Albrecht et al, 2021, Cadigan et al, 2012). Importantly, Wnt-β-catenin signaling is frequently upregulated in human cancer, particularly in colorectal cancer, where mutations in adenomatous polyposis coli (APC) cause hyper-stabilization of β-catenin (Schneikert et al, 2007, Zhan et al, 2017). Consequently, Wnt target genes including genes regulating the cell cycle are aberrantly expressed, thereby contributing to tumorigenesis and tumor progression (He et al, 1998, Tetsu et al, 1999). Wnt signaling can also stabilize proteins other than β-catenin. This is referred to as the “Wnt Stabilization Of Proteins (Wnt/STOP)” pathway, which was shown to be involved in the regulation of cell size and mitosis (Acebron et al, 2014, Hinze et al, 2019, Huang et al, 2015, Stolz et al, 2015b, Taelman et al, 2010). In fact, inhibition of β-catenin-independent Wnt signaling causes chromosome missegregation in mitosis leading to aneuploidy in human somatic cells as well as in pluripotent stem cells (Augustin et al, 2017, Jaime-Soguero d et al, 2024, Lin et al, 2021, Stolz et al, 2015b).

Perpetual mitotic chromosome missegregation promotes the evolvement of aneuploidy, which represent the basis for whole chromosome instability (W-CIN), a major form of genome instability and a hallmark of human cancer (Bakhoum et al, 2017, Chen X et al, 2024, Hanahan D & Weinberg RA, 2011, Lukow et al, 2022, Sansregret et al, 2018). W-CIN originates during mitosis and can be driven by various abnormalities leading to chromosome missegregation (Bastians & H., 2015, Devillers et al, 2024). Specifically, an abnormal increase in growth rates of microtubules within the mitotic spindle, which impairs the proper positioning of the spindle has been recently identified as an important mechanism leading to W-CIN in human cancer cells (Ertych et al, 2014, Ertych et al, 2016, Paim et al, 2024, Schmidt et al, 2021, Tamura et al, 2020). In fact, chromosomally unstable cancer cells that are characterized by high levels of chromosome missegregation typically display increased spindle microtubule growth rates as a driver for aneuploidy, and thus, for W-CIN (Bohly et al, 2022, Ertych et al, 2014, Tamura et al, 2020).

In contrast to W-CIN, structural CIN (S-CIN) causes structural chromosome aberrations including DNA amplifications, deletions and translocations (Siri et al, 2021). S-CIN is often associated with chromosomal breaks that are driven by defects in DNA repair or by slowed or stalled DNA replication, a condition known as DNA replication stress (Siri et al, 2021, Zeman et al, 2014). Overall, CIN represents a major form of genome instability and is a hallmark of human cancer (Hanahan D & Weinberg RA, 2011, Sansregret et al, 2018). Consequently, CIN promotes tumor evolution by generating genomic heterogeneity in cancer cells, thereby supporting aggressive growth phenotypes, metastasis and therapy resistance (Bakhoum et al, 2017, Chen X et al, 2024, Lukow et al, 2022, Sansregret et al, 2018).

Interestingly, S-CIN and W-CIN are typically detected concomitantly in chromosomally unstable cancer cells suggesting functional links between the two forms of CIN (Burrell RA et al, 2013, Janssen et al, 2011, Tijhuis et al, 2019). Indeed, several studies showed that replication stress can not only trigger S-CIN, but also affects mitosis leading to chromosome missegregation and this can be mediated by increase microtubule dynamics (Bohly et al, 2019, Bohly et al, 2022, Burrell RA et al, 2013, Dwivedi et al, 2023, Wilhelm et al, 2019). In addition, we recently reported that, in pluripotent stem cells, loss of Wnt can impact on DNA replication and, thereby supporting the generation of mitotic errors (Jaime-Soguero d et al, 2024). However, it remains unclear what mechanisms and pathways contribute to abnormal microtubule dynamics as a trigger for chromosome missegregation in response to replication stress. Here, we show that in human cancer cells Wnt10b signaling is required upon replication stress to ensure normal microtubule dynamics from S phase until mitosis and to prevent chromosomal breaks and mitotic errors without affecting DNA replication dynamics *per se*. Hence, we propose that Wnt10b signaling acts in cancer cells during S phase-associated replication stress as a rescue pathway to limit genome instability.

## Results

### Loss of Wnt10b signaling causes mitotic errors

We have previously shown that loss of Wnt10b signaling causes mitotic errors in human somatic cells (Fig. 1A). Of note, inhibition of Wnt(10b) signaling in chromosomally stable colorectal cancer cells (HCT116), either at the canonical (co)receptor level by DKK1 treatment, by knockout of the Wnt secretion factor *EVI/WNTLESS* (Augustin et al, 2017) or of *WNT10B* ligand (Fig. S1A) triggered increased microtubule growth rates in mitotic cells as determined by live cell microscopy tracking of individual microtubule plus tips within mitotic spindles (Fig. 1B,C). Increased mitotic microtubule growth rates, in turn, were associated with increased chromosome missegregation during mitosis as detected by the enhanced occurance of lagging chromosomes during anaphase of mitosis (Fig. 1D,E). The causal relationship between increased microtubule growth rates and chromosome missegregation was validated by direct suppression of chromosome missegregation after restoration of proper microtubule growth rates using sub-nanomolar concentrations of the microtubule-binding drug Taxol as shown in previous studies (Fig. 1C,E)(Ertych et al, 2014, Stolz et al, 2015b). In all conditions of suppressed Wnt signaling the mitotic errors were suppressed upon GSK3β kinase inhibition indicating that Wnt-GSK3β signaling is involved in mitotic regulation (Fig. 1C,E)(Jaime-Soguero d et al, 2024, Lin et al, 2021). Importantly, only treatment with purified Wnt10b, but not with Wnt3a protein, rescued both, abnormally increased mitotic microtubule growth rates and chromosome missegregation in HCT116-*WNT10B* and HCT116-*EVI/WNTLESS* knock-out cells, even though Wnt3a was much more potent to activate Wnt/β-catenin signaling (Fig. 1C,E, Fig. S1B) supporting the notion that β-catenin independent Wnt10b signaling is required for faithful chromosome segregation (Lin et al, 2021, Stolz et al, 2015b).

**Figure 1.**
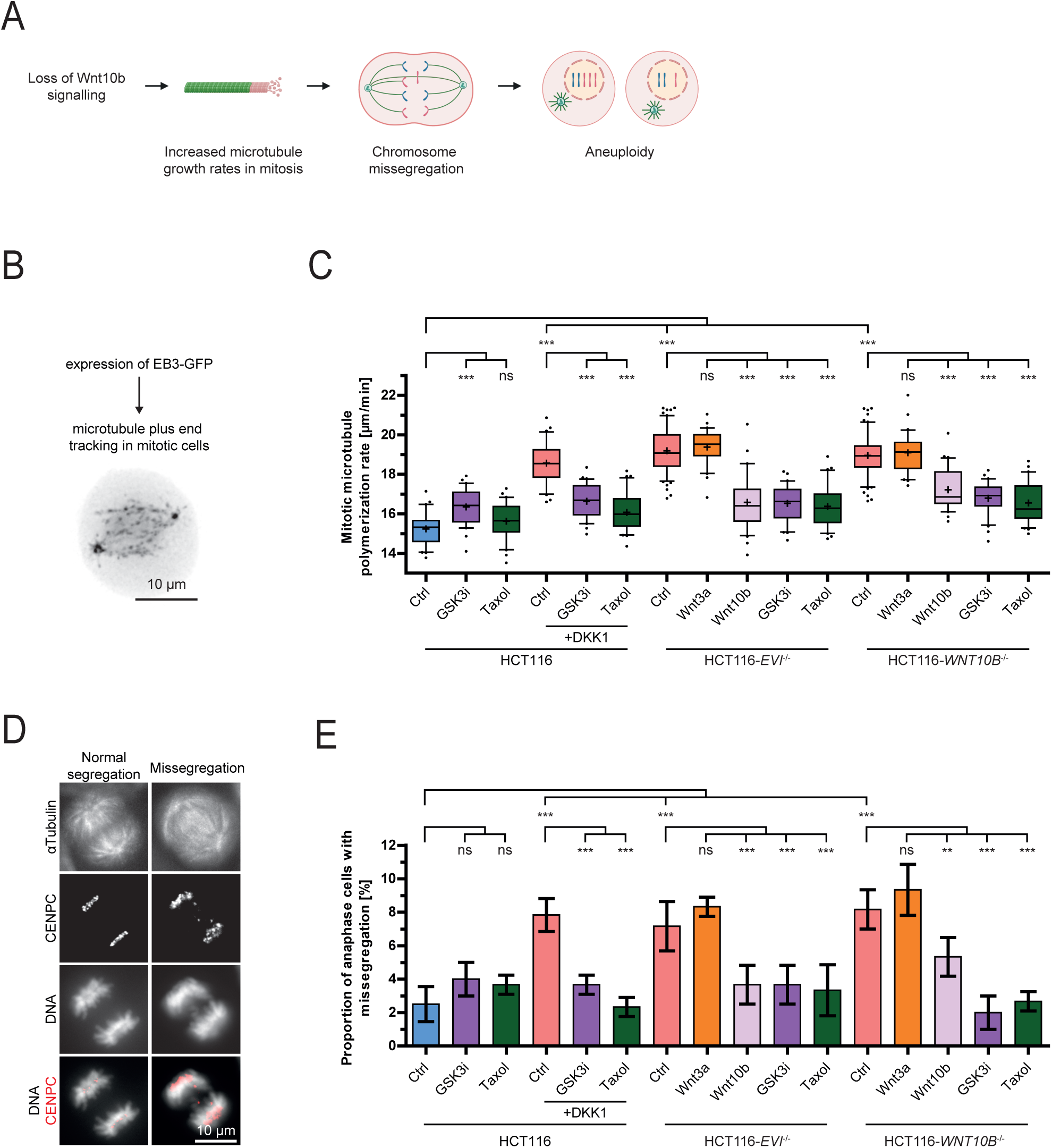
Wnt10b signaling is required for normal microtubule dynamics and faithful chromosome segregation during mitosis. **(A)** Model depicting the relationship between the loss of Wnt signaling, increased mitotic microtubule plus-end growth rates, chromosome missegregation and the induction of aneuploidy. **(B)** Experimental outline for the measurement of microtubule plus-end growth rates. **(C)** Measurements of mitotic microtubule growth rates in HCT116, HCT116-*EVI* and HCT116-*WNT10B* knock-out cells after Wnt inhibition. As indicated, cells were treated with DMSO (Ctrl), 600 ng/ml DKK1, 0.6 µM GSK3 inhibitor (CHIR-99021), 0.2 nM Taxol or 400 ng/ml of recombinant Wnt ligands. Measurements are based on analysis of 25 microtubules/cell (n = 30 cells from three independent experiments, two-tailed *t-*test). **(D)** Example images of cells with and without mitotic chromosome missegregation. Fixed cells were stained for spindle microtubules (α-tubulin), kinetochores (Cenp-C and chromosomes (DNA) to detect lagging chromatids during anaphase of mitosis, scale bar 10 µm. **(E)** Quantification of the proportion of cells exhibiting chromosome missegregation upon Wnt inhibition. Cells were treated as described in (C) and anaphase cells with lagging chromosomes were quantified. Graphs show mean values ± SD (n = 300 cells from three independent experiments, two-tailed *t*-test).

### Wnt10b signaling is required during S phase to ensure faithful mitotic chromosome segregation

Since Wnt10b signaling is required for proper mitosis we expected it to act immediately before or during mitosis to ensure normal mitotic microtubule dynamics and faithful chromosome segregation. To test this, we treated cell cycle-synchronized HCT116 cells with DKK1 to induce mitotic errors or synchronized HCT116-*EVI/WNTLESS* knock-out cells with purified Wnt10b to rescue mitotic errors at specified phases of the cell cycle (Fig. 2A, Fig. S2). Surprisingly, DKK1-mediated inhibition of Wnt signaling for only two hours during S phase, but not during G2 phase or immediately before or during mitosis (G2/M) induced GSK3β-dependent abnormal microtubule growth rates and chromosome missegregation (Fig. 2B, 2C). *Vice versa*, treatment of HCT116-*EVI/WNTLESS* knockout cells with purified Wnt10b ligand rescued mitotic errors only when applied for two hours during S phase, but not during later stages of the cell cycle, whereas treatment with Wnt3a from S phase until mitosis had no effect (Fig. 2D,E). Thus, Wnt10b signaling seems to be required specifically during S phase to ensure faithful chromosome segregation during the subsequent mitosis. To extend these findings we used chromosomally unstable (CIN+) colorectal cancer cells, which are characterized by inherently high rates of chromosome missegregation that are caused by increased microtubule growth rates (Bohly et al, 2022, Ertych et al, 2014, Tamura et al, 2020). Also, in these CIN+ cancer cells treatment with Wnt10b, but not Wnt3a specifically during S phase efficiently rescued abnormal microtubule growth rates and chromosome missegregation in mitosis (Fig. 2F,G) indicating that Wnt10b signaling is important during S phase to suppress mitotic errors in human cancer cells.

**Figure 2.**
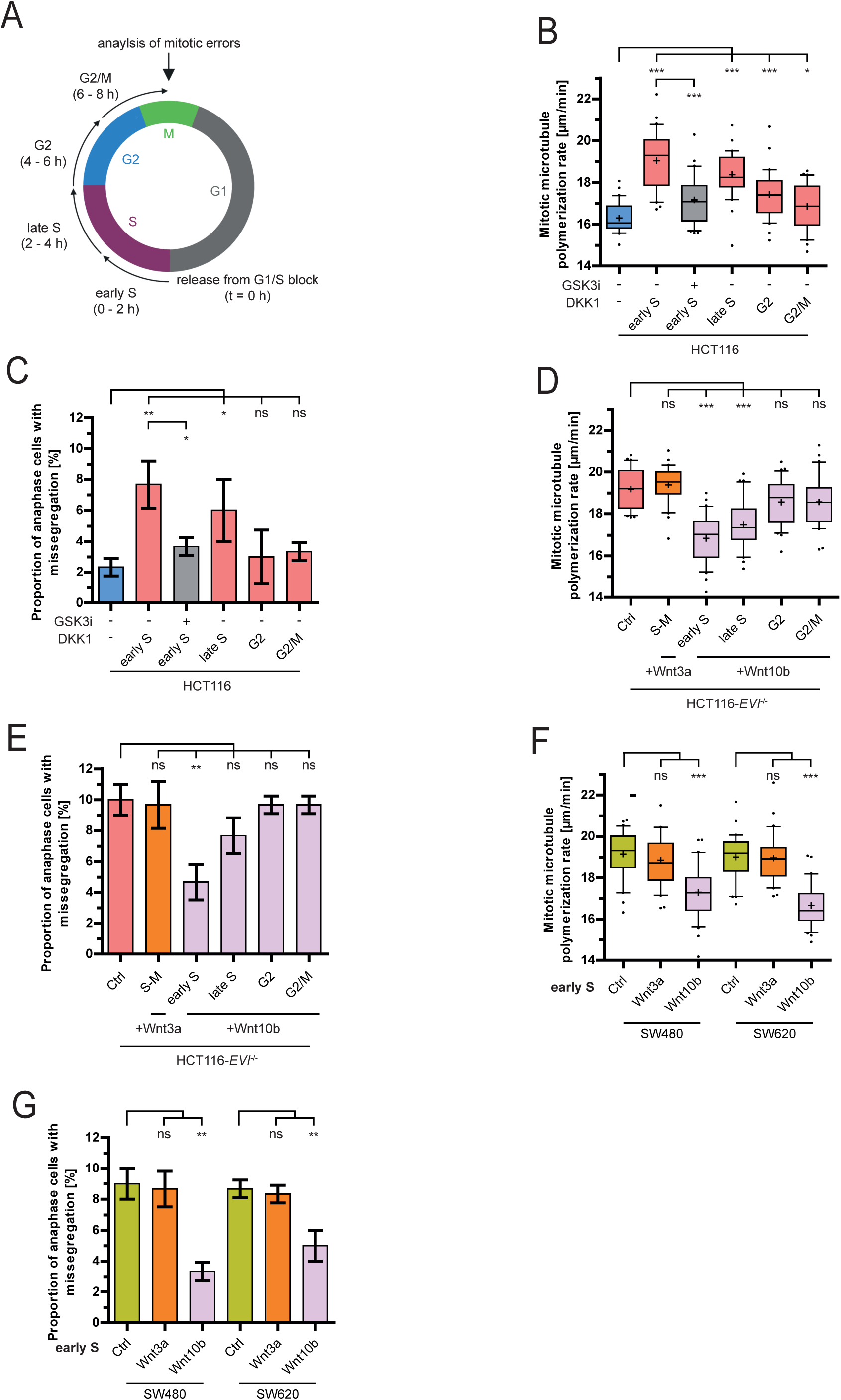
Wnt10b acts during S phase to promote faithful mitosis. **(A)** Schematic outlining cell cycle specific Wnt modulation using cell cycle synchronized cells followed by analysis of mitotic outcomes. **(B)** Measurements of microtubule growth rates in HCT116 cells after cell cycle phase-specific treatment with 600 ng/ml DKK1 in the absence or presence of 0.6 µM GSK3β kinase inhibitor (25 microtubules/cell, n = 30 cells from three independent experiments, two-tailed *t*-test). **(C)** Quantification of anaphase cells showing chromosome missegregation upon treatments as described in (B). Graph shows mean values ± SD (n = 300 cells from three independent experiments, two-tailed *t*-test). **(D)** Measurements of microtubule growth rates in HCT116-*EVI/WNTLESS* knock-out cells after cell cycle-specific treatment with 400 ng/ml of recombinant Wnt ligands (25 microtubules/cell, n = 30 cells from three independent experiments, two-tailed *t*-test). **(E)** Quantification of HCT116-*EVI/WNTLESS* knock-out cells showing chromosome missegregation after treatments as described in (D) (mean ± SD, n = 300 cells from three independent experiments, two-tailed *t*-test). **(F)** Measurements of microtubule growth rates in chromosomally unstable (CIN+) SW480 and SW620 colorectal cancer cells with or without treatment with 400 ng/ml Wnt ligands during early S phase for two hours (25 microtubules/cell, n = 30 cells from three independent experiments, two-tailed *t*-test). **(G)** Quantification of CIN+ cancer cells displaying chromosome missegregation with or without treatments as in (F) (mean ± SD, n = 300 cells from three independent experiments, two-tailed *t*-test).

### Inhibition of Wnt signaling increases microtubule growth rates from S phase into mitosis

Since Wnt10b signaling is required during S phase but regulates mitosis several hours later we wondered whether inhibition of Wnt signaling impacts microtubule behavior already before cells enter mitosis. Thus, we determined interphase microtubule growth rates in synchronized cells with or without Wnt inhibition (Fig. 3A). Interestingly, DKK1 treatment or loss of *EVI/WNTLESS* resulted in increased microtubule growth rates in S phase and this abnormality was maintained until mitosis (Fig. 3B). The same findings were obtained using synchronized CIN+ SW480 and SW620 cells that exhibited inherently high microtubule polymerization rates from S phase until mitosis (Fig. 3C). Importantly, treatment of synchronized Wnt-inhibited or CIN+ cancer cells with low doses of Taxol just before mitosis (Fig. 3D) was sufficient to restore abnormal microtubule growth rates and suppressed chromosome missegregation (Fig. 3E,F) indicating that the abnormal microtubule growth rates originate in S phase, and they are maintained until mitosis as a “cytoskeletal memory”, which ultimately causes missegregation of chromosomes during mitosis.

**Figure 3.**
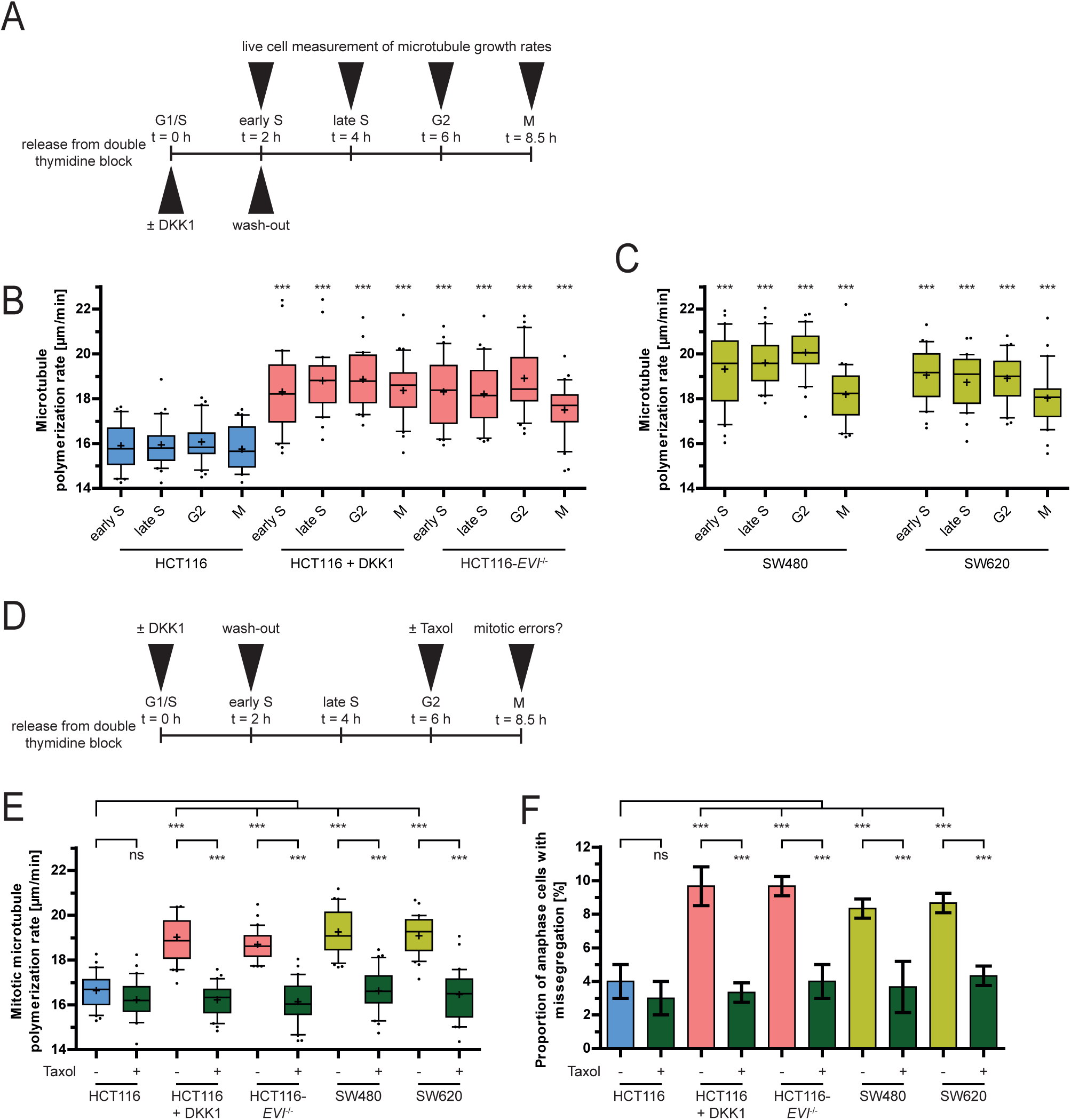
Wnt inhibition causes increased microtubule growth rates from S phase until mitosis. **(A)** Experimental setup for the cell cycle-specific analysis of microtubule growth rates. **(B)** Cell cycle stage dependent measurements of microtubule growth rates after inhibition of Wnt signaling in HCT116 upon DKK1 treatment or in HCT116 *EVI/WNTLESS* knock-out cells (20 microtubules/cell, n = 30 cells from three independent experiments). **(C)** Cell cycle stage dependent measurements of microtubule growth rates in chromosomally unstable colorectal cancer cells (20 microtubules/cell, n = 30 cells from three independent experiments). **(D)** Experimental setup for the analysis of mitotic microtubule assembly rates upon 0.2 nM Taxol treatment. **(E)** Measurements of mitotic microtubule growth rates in the indicated cells with or without Taxol treatment at G2/M (20 microtubules/cell, n = 30 cells from three independent experiments). **(F)** Quantification of cells showing chromosome missegregation after treatment as in (E)(mean ± SD, n = 300 cells from three independent experiments, two-tailed *t*-test).

### Replication stress does not link Wnt signaling and chromosome missegregation

Since Wnt inhibition acts specifically during S phase to regulate mitosis we wondered whether Wnt10b signaling might be linked to DNA replication, which is the main process during S phase of the cell cycle. In fact, it is well established that DNA replication stress is a frequent condition in chromosomally unstable cancer cells and associated with broad genome instability (Gaillard et al, 2015, Zeman et al, 2014). Moreover, recent studies have highlighted that replication stress also impacts mitosis and causes abnormal microtubule dynamics, thereby explaining the concomitant presence of replication stress and high chromosome missegregation rates in chromosomally unstable cancer cells (Bohly et al, 2019, Bohly et al, 2022, Burrell et al, 2013, Fragkos et al, 2017, Liu et al, 2014, Wilhelm et al, 2020). In agreement with this, we found that induction of replication stress in chromosomally stable HCT116 cells by treatment with low concentrations of aphidicolin, a specific inhibitor of DNA polymerases and well-established inducer of replication stress (Bohly et al, 2019, Bohly et al, 2022, Ikegami et al, 1978, Minocherhomji et al, 2015), increased mitotic microtubule growth rates and chromosome missegregation (Fig. 4A,B). *Vice versa*, increased microtubule growth rates and chromosome missegregation endogenously present in CIN+ cancer cells were suppressed upon nucleoside supplementation, an established mean to alleviate replication stress (Bohly et al, 2019, Burrell et al, 2013, Wilhelm et al, 2014) (Fig. 4C,D). Thus, replication stress during S phase mimics Wnt inhibition with respect to the induction of mitotic errors raising the question whether Wnt10b inhibition causes replication stress. To address this, we employed DNA combing analysis, the gold standard method to measure the progression of individual replication forks during DNA replication (Fig. 4E)(Moore et al, 2022). As expected, treatment with aphidicolin reduced replication fork velocities (Fig. 4F). Similarly, CIN+ cancer cells exhibited slowed fork progression indicating endogenous replication stress in chromosomally unstable cancer cells as reported previously (Fig. 4F)(Bohly et al, 2019, Bohly et al, 2022, Burrell et al, 2013). However, neither DKK1 treatment nor the loss of *EVI/WNTLESS* (HCT116-*EVI*/*WNTLESS* knock-out cells) affected replication fork speed (Fig. 4F). Accordingly, no reduced inter-origin distances were found upon Wnt inhibition (Fig. S3A), which is known to be a direct consequence of replication stress (Bohly et al, 2022, Ibarra et al, 2008, Moiseeva et al, 2019). We also did not find a delay in S phase progression, which would be otherwise expected in the presence of replication stress (Bartkova et al, 2005, Gorgoulis et al, 2005) (Fig. S3C,D). Also, treatment of CIN+ cells with Wnt10b, which rescued mitotic errors (Fig. 2F,G), did not affect replication stress in these cancer cells (Fig. 4F). Together, inhibition of Wnt10b signaling affects mitotic chromosome segregation, but does not directly impact on DNA replication dynamics in human cancer cells.

**Figure 4.**
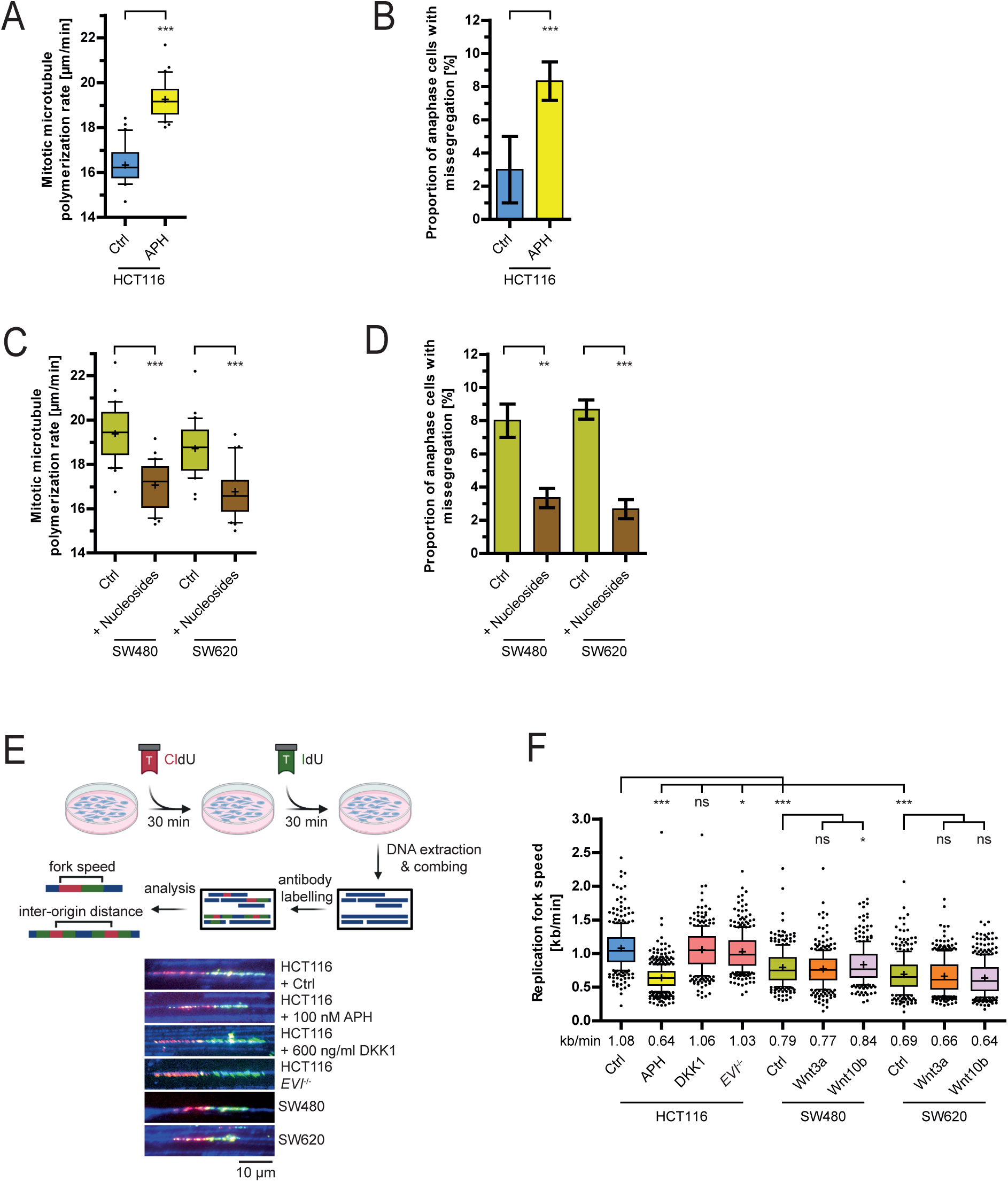
Replication stress does not link Wnt signaling and chromosome missegregation. **(A)** Measurement of microtubule polymerization rates in HCT116 cells after induction of replication stress. Cells were treated with 100 nM aphidicolin for 16 hours prior to live cell measurements of microtubule growth rates (25 microtubules/cell, n = 30 cells from three independent experiments, two-tailed *t*-test). **(B)** Quantification of anaphase cells showing chromosome missegregation upon replication stress. HCT116 cells were treated as in (A) and chromosome missegregation was detected in anaphase cells (mean ± SD, n = 300 cells from three independent experiments, two-tailed *t*-test). **(C)** Measurement of mitotic microtubule growth rates in CIN+ cancer cells upon alleviation of replication stress by nucleoside supplementation (25 microtubules/cell, n = 30 cells from three independent experiments, two-tailed *t*-test). **(D)** Quantification of anaphase cells showing chromosome missegregation upon treatment as in (C) (mean ± SD, n = 300 cells from three independent experiments, two-tailed *t*-test). **(E)** Schematic setup for DNA combing to determine replication fork speed and inter-origin distances. Representative examples of labelled unidirectional DNA fibers are shown. **(F)** Determination of replication fork progression rates using the indicated cells and treatments (>300 fibers per condition, n=1, two-tailed *t*-test).

### Wnt10b signaling acts downstream of replication stress to ensure faithful mitosis

Since Wnt inhibition does not impact on replication dynamics *per se*, we hypothesized that Wnt10b signaling might act downstream of replication stress to ensure proper mitosis. This hypothesis was supported by the rescue of mitotic errors in CIN+ cells, whose mitotic errors are triggered by endogenous replication stress (Fig. 4C,D,F), after the activation Wnt10b signaling (Fig. 2 F,G). To further address this, we induced replication stress during early S phase by pulse-treatment with aphidicolin or hydroxyurea, which depletes cellular nucleotide pools (Fig. 5A)(Bianchi et al, 1986). Both means of replication stress induction resulted in increased microtubule growth rates and chromosome missegregation in the subsequent mitosis, both of which were suppressed by co-treatment with Wnt10b, but not with Wnt3a (Fig 5B,C). Since replication stress-induced abnormal microtubule growth rates may originate from S phase (Fig. 3) we also determined microtubule growth rates during S phase upon induced replication stress (Fig. 5A) or using CIN+ cancer cells, which suffer from endogenous replication stress (Fig. 4F). In fact, replication stress induced increased microtubule growth rates already in S phase where they were suppressed by Wnt10b treatment (Fig. 5D). Together, our data indicate that Wnt10b signaling can act as a pathway during S phase that suppresses mitotic errors induced by DNA replication stress in human cancer cells.

**Figure 5.**
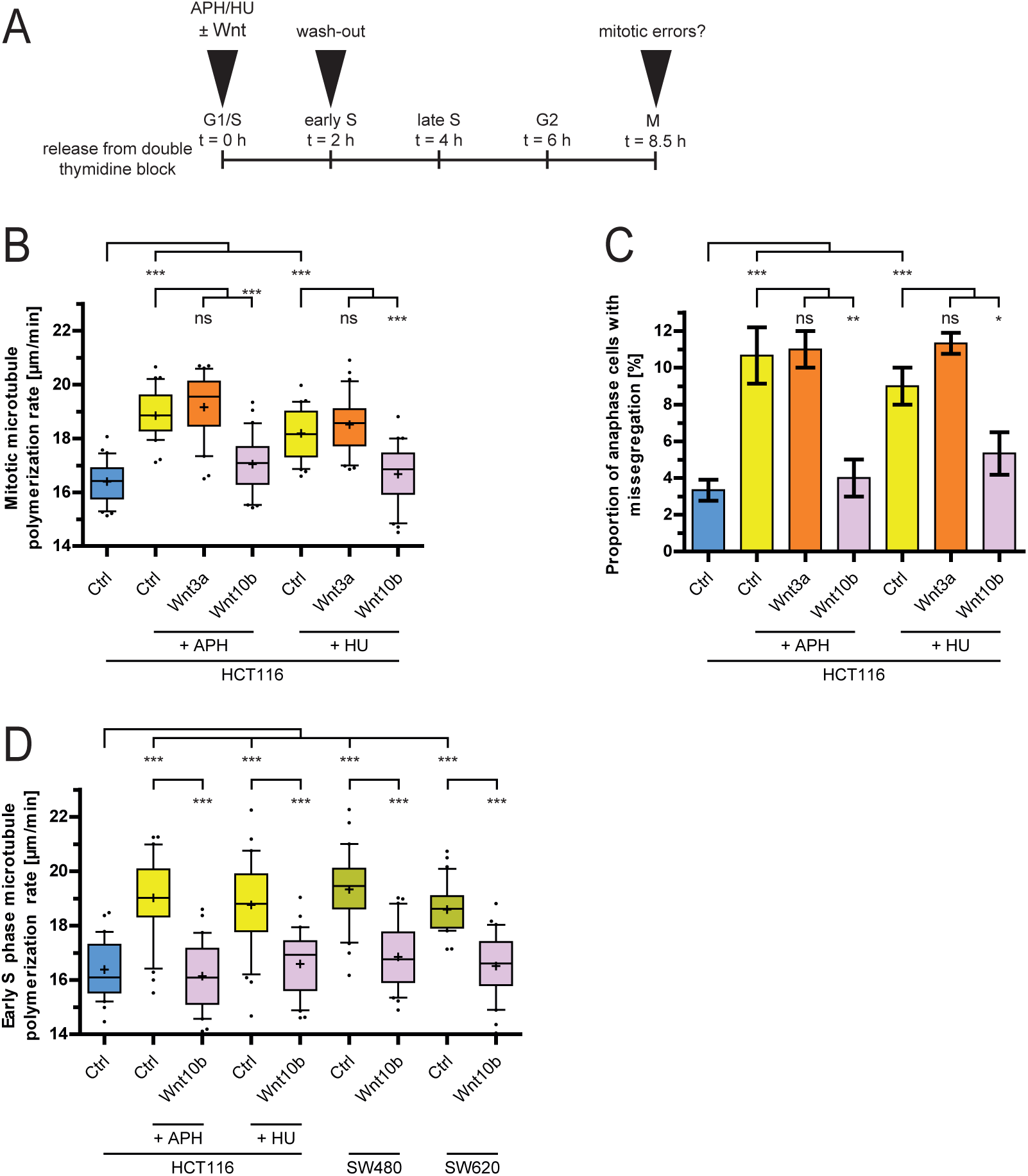
Wnt10b signaling ensures faithful chromosome segregation after replication stress. **(A)** Experimental outline for the analysis of mitotic errors after replication stress and Wnt treatment. **(B)** Measurement of mitotic microtubule growth rates in HCT116 cells after treatment with 100 nM aphidicolin (APH) or 2 mM hydroxyurea (HU) in the presence or absence of 400ng/ml Wnt ligands (25 microtubules/cell, n = 30 cells from three independent experiments, two-tailed *t*-test) **(C)** Quantification of anaphase cells showing chromosome missegregation after treatment as in (B) (mean ± SD, n = 300 cells from three independent experiments, two-tailed *t*-test). **(D)** Measurements of microtubule growth rates in early S phase upon replication stress. HCT116, SW480 and SW620 cells were synchronized in early S phase, treated as indicated and microtubule growth rates were analyzed during S phase (25 microtubules/cell, n = 30 cells from three independent experiments, two-tailed *t*-test).

### Wnt10b signaling suppresses replication stress-induced chromosome breaks

In addition to causing mitotic errors, replication stress is known to give rise to chromosomal breaks, which form the basis of structural chromosome instability in cancer (Branzei et al, 2010, Zeman et al, 2014). Since Wnt10b regulates mitotic errors in response to replication stress, we wondered if Wnt10b signaling also affects the downstream generation of chromosomal breaks. To address this, we induced replication stress with aphidicolin or hydroxyurea treatment during early S phase, either in the presence or absence of Wnt10b or Wnt3a (Fig. 6A). As expected, we observed increased chromosomal breakage upon replication stress as determined by chromosome spread analysis (Fig. 6B). Intriguingly, treatment with Wnt10b, but not with Wnt3a, significantly suppressed replication stress-induced chromosomal breaks (Fig. 6C), indicating that Wnt10b indeed modulates the generation of chromosomal breaks upon replications stress. Similarly, increased levels of chromosomal breaks were also detectable in CIN+ cancer cells, which were suppressed upon alleviation of replication stress upon nucleoside supplementation, indicating that these chromosomal breaks resulted from endogenous replication stress present in these cancer cells (Fig. 6D). Importantly, the replication stress-induced chromosomal breaks in these CIN+ cancer cells were also efficiently suppressed upon Wnt10b, but not Wnt3a treatment (Fig. 6D). Further supporting the role of Wnt10b signaling in suppressing chromosomal breaks downstream of replication stress, we found that inhibition of Wnt signaling either upon DKK1 treatment or in *EVI/WNTLESS* knockout cells induced Wnt10b-dependent chromosomal breaks as seen after replication stress (Fig. 6C). Taken together, our results indicate that Wnt10b signaling functions as a rescue pathway downstream of replication stress, preventing both mitotic chromosome missegregation as well as chromosome breaks, two well-established consequences of DNA replication stress and hallmarks of genome instability in human cancer.

**Figure 6.**
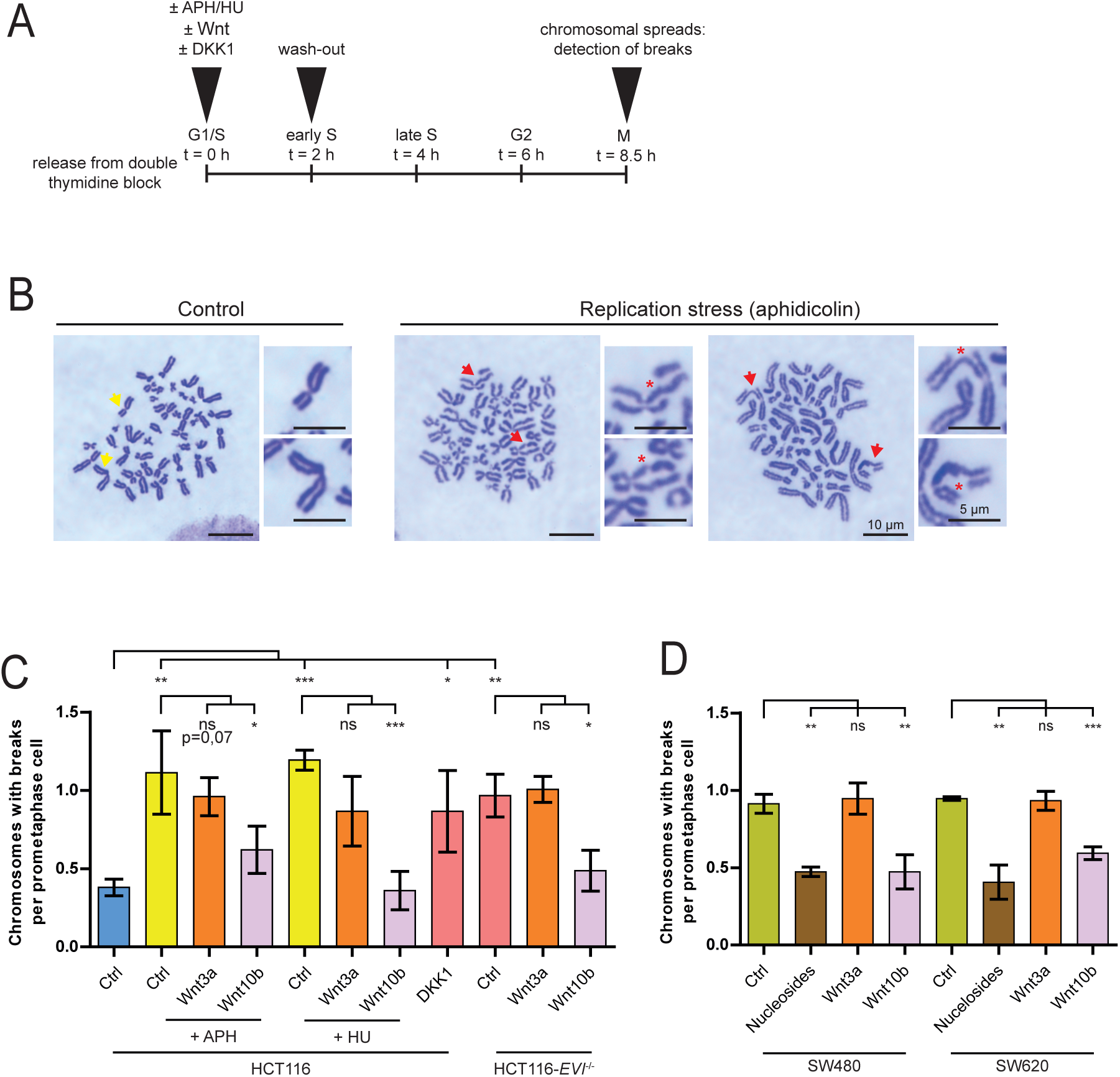
Wnt10b suppresses replication stress-induced chromosomal breaks. **(A)** Setup for the analysis of chromosomal breaks. **(B)** Representative images of chromosome spreads with or without replication stress-induced chromosomal breaks. Scale bar, 5 µm and 10 µm as indicated. **(C)** Quantification of chromosome breaks after replication stress or Wnt inhibition. Mitotic chromosome spreads were analyzed from cells after induction of replication stress (100 nM APH or 2mM HU) or after Wnt inhibition (DKK1, *EVI/WNTLESS* knockout) in the presence or absence of Wnt ligands. The graph shows the number of chromosomes with breaks per cell (mean ± SD, n=150 chromosome spreads from 3 experiments; two-tailed *t*-test). **(D)** Quantification of chromosome breaks in CIN+ cancer cells after treatment with Wnt ligands or upon nucleosite supplementation. The graph shows the number of chromosomes with breaks per cell (mean ± SD, n=150 chromosome spreads from 3 experiments; two-tailed *t*-test).

## Discussion

Our work revealed a yet unrecognized und unexpected function of Wnt10b signaling to protect cells from mitotic errors and chromosomal breaks in response to DNA replication stress, which is a major source for chromosomal instability (CIN) in human cancer (Igarashi et al, 2024, Zeman et al, 2014). Thus, Wnt10b signaling might act as a suppressor of CIN that is a well-established driving force for the generation of high genetic heterogeneity and variability in cancer supporting tumor evolution towards aggressive growth phenotypes, metastasis and therapy resistance (Chen et al, 2025, Sansregret et al, 2018).

Replication stress (RS) is a highly cancer-relevant condition that is characterized by slowed or stalled DNA replication during S phase of the cell cycle. It can be triggered by various cellular conditions including nucleotide shortage, the presence of abnormal chromosome structures, DNA damage or activation of oncogenes such as *MYC, RAS* or *CYCLIN-E* and contributes to the induction of structural chromosome aberrations that are frequently seen in cancer (Igarashi et al, 2024, Zeman et al, 2014). In addition, recent work has uncovered that RS also affects mitosis by deregulating mitotic microtubule dynamics and centrosome separation leading to chromosome missegregation, thereby driving the evolvement of aneuploidy (Bohly et al, 2019, Bohly et al, 2022, Burrell et al, 2013, Jaime-Soguero d et al, 2024, Wilhelm et al, 2019). With this, RS sits at the helm of CIN in cancer cells.

Intriguingly, our work demonstrates that activation of Wnt10b signaling in cancer cells suffering from RS is sufficient to suppress both, the generation of chromosomal breaks and of mitotic errors indicating that Wnt10b acts to limit deleterious consequences of RS in cancer cells. In line with a function of Wnt10b downstream of RS, we showed that Wnt10b signaling does not impact on DNA replication dynamics in cancer cells *per se* and its inhibition neither causes slowed replication fork progression nor activation of dormant origin firing, which is a cellular rescue mechanism to compensate for slow replication and to ensure completion of DNA replication (Bohly et al, 2022, Ibarra et al, 2008, Moiseeva et al, 2019). However, this stands in contrast to our recent findings in pluripotent stem cells where Wnt3a signaling suppresses origin firing in S phase (Jaime-Soguero d et al, 2024), which is also required for faithful mitotic chromosome missegregation (Bohly et al, 2022, Jaime-Soguero d et al, 2024). It is note that pluripotent stem cells have high basal levels of replicative stress and unique replication profiles (Kafer et al, 2022, Kurashima et al, 2024), which may contribute to the difference on the Wnt involvement in replicative stress in comparision to cancer cells. Further analysis on proteins that are regulated by Wnt3a and Wnt10b during S phase in stem cells and somatic cells, respectively, are required to define the overlapping or non-overlapping pathways involved in S phase and mitotic regulation in both cell systems.

Cancer cells typically exhibit mild RS, which is sufficient to trigger low levels of structural chromosome aberrations and mitotic errors that are still compatible with cell survival and avoiding cell cycle arrest. Severe RS, however, causes excessive stalling of replication forks often associated with severe DNA damage, which reduces cell viability and causes cell cycle arrest via the activation of replication checkpoint pathways (Zeman et al, 2014). These checkpoint mechanisms involving kinases such as ATR and Chk1 fulfills protective functions to replication stressed cells by imposing cell cycle halt and ensuring efficient re-start of fork progression upon repair and thereby, these checkpoint functions are essential to cell viability (Simoneau et al, 2021). In contrast to checkpoint activation, Wnt10b acts downstream of mild replication stress in the absence of checkpoint activation either induced by very low concentrations of aphidicolin or endogenously present in chromosomally unstable colorectal cancer cells (Bohly et al, 2019, Burrell et al, 2013). To avoid deleterious consequences of RS, cancer cells evolved various adaptation mechanisms to cope with RS. For instance, upregulation of proteins such as Timeless and Claspin is observed in cancer cells suffering from replication stress and this alleviates slowed fork progression independently of the ATR-Chk1 checkpoint (Bianco et al, 2019). However, as shown by our DNA combing analysis, Wnt10b addition does not improve replication fork velocities in cancer cells with mild RS indicating that Wnt10b does not act directly on replication fork progression. Nevertheless, Wnt10b clearly has protective functions in cells with RS and hence, upregulation of Wnt10b might be useful for cells suffering from RS. In fact, in many cancer types Wnt10b is found to be upregulated, but it is currently unclear whether this is associated with RS in these tumors (Perkins et al, 2023).

It is yet unknown how Wnt10b signaling suppresses mitotic errors and chromosomal breaks in response to RS. It can be speculated that the alterations in microtubule dynamics seen upon Wnt10b modulation are involved. Our work provides new evidence that abnormally increased microtubule growth rates are an immediate response to RS readily during S phase and this alteration is maintained until cells enter mitosis where it causes chromosome missegregation by affecting proper spindle positioning as shown previously (Ertych et al, 2014). Thus, RS or Wnt inhibition triggers a rapid increase of microtubule growth rates, which might have a yet unknown physiological function in S phase. Interestingly, recent reports have indicated that dynamic microtubules are involved in nuclear DNA repair through regulation of chromatin mobility involving the microtubule-LINC (linker of nucleoskeleton and cytoskeleton) complex (Lawrence et al, 2016, Lottersberger et al, 2015). Such microtubule-involving repair mechanisms might be activated upon RS to prevent additional effects during DNA replication, but further detailed investigations are needed to address this intriguing possibility.

It is remarkable that specifically Wnt10b acts during S phase of the cell cycle although it has been suggested to activate canonical Wnt signaling involving Frizzled receptors, LRP co-receptors, GSK3β kinase and the well-characterized β-catenin destruction complex (Wend et al, 2012). However, we found that Wnt3a is much more potent to activate β-catenin dependent Wnt signaling while have no function in response to RS indicating a highly specialized and β-catenin independent role of Wnt10b during S phase. In agreement, we previously demonstrated that LRP and GSK3β-dependent, but β-catenin-independent Wnt/STOP (Wnt mediated stabilization of proteins) signaling regulates microtubule dynamics causing mitotic chromosome missegregation and aneuploidy (Acebron et al, 2014, Acebron et al, 2016, Huang et al, 2015, Stolz et al, 2015a, Stolz et al, 2015b). Based on this, we suggest that Wnt10b-activated signaling, although employing the same Wnt pathway components, specifically regulates target proteins, possibly through Wnt/STOP, during S phase to contribute to its function upon RS.

Overall, our work highlights Wnt10b signaling as an extracellular signaling pathway regulating consequences of DNA replication stress and mitotic errors in human cancer cells. In the same line, we recently reported similar roles for other extracellular signaling pathways such as BMP, WNT and FGF signaling in embryonic stem cells linking genome instability to cell fate (Jaime-Soguero d et al, 2024) suggesting key roles for extracellular cues in the maintenance of genome stability.

## Materials and Methods

### Cell culture

HCT116 (RRID:CVCL_0291), SW480 (RRID:CVCL_054), SW620 (RRID:CVCL_0547) and HEK293T (RRID:CVCL_0063) cells were purchased from ATCC (Manassas, USA) and cultured in RPM1-1640 or DMEM medium (PAN Biotech, Germany). HCT116-*EVI/WNTLESS* knock-out cells were described previously (Augustin et al, 2017, Lin et al, 2021). Media were supplemented with 10% FBS (#AC-AB-0024, Anprotec, Germany) and 1% Penicillin/streptomycin (#AC-AB-0024, Anprotec, Germany) in a CO_2_ incubator at 37°C with 5% CO2.

### Generation of *WNT10B* knockout cells

HCT116-*WNT10B* knock-out cells were generated as previously described (Ran et al, 2013, Voloshanenko et al, 2017). A short-guide RNA for CRISPR/Cas9 (3’-GGAAGAATGCGGCTCTGACA-5’) was designed using E-CRISP (http://www.e-crisp.org), purchased from Eurofins Inc. (Ebersberg, Germany) and cloned into a pSpCas9(BB)-2A-Puro (px459) plasmid, which was a gift from Feng Zhang (Addgene plasmid #48139; http://n2t.net/addgene:48139; RRID:Addgene_48139)(Ran et al, 2013). HCT116 cells were transfected and selected in culture medium containing 2 µg/ml of puromycin. Pools of cells were first expanded and a single cell clone was generated and further analyzed.

### Cell treatments

Where indicated, cells were treated with 100 nM aphidicolin (sc-201535, Santa Cruz, USA) or 2 mM hydroxyurea (H8627-1G, Merck-Millipore, Burlington, UK) to induce replication stress. Replication stress was alleviated by treatment with additional nucleosides for 48 h (30 μM 2ʹ-deoxyadenosine monohydrate (#sc-216290, Santa Cruz, USA), 30 μM 2ʹ-deoxycitidine hydrochloride (#sc-220820, Santa Cruz, USA), 30 μM thymidine (#sc-296542, Santa Cruz, USA) and 30 μM 2ʹ-deoxyguanosine mono-hydrate (#sc-238433, Santa Cruz, USA) as described previously (Bohly et al, 2019, Wilhelm et al, 2014). Cells were treated with 0.6 µM CHIR99021 (SML1046, Sigma Aldrich, Germany) to inhibit GSK3β kinase. Increased microtubule growth rates were suppressed by treatment with 0.2 nM Taxol (Paclitaxel, T7191-1MG, Sigma Aldrich, Germany) as shown previously (Ertych et al, 2014). As controls cells were treated with H_2_O or dimethyl sulfoxide (D1418, Sigma-Aldrich, Germany). Cells were treated with 400 ng/ml of human recombinant Wnt3a (5036-WN/CF, R&D Systems, Germany) or human recombinant Wnt10b (7196-WN/CF, R&D Systems, Germany) to induce ligand-specific Wnt signaling, or with 600 ng/ml of human recombinant DKK1 (120-30-500, PeproTech, Germany) to inhibit Wnt/LRP6 signaling. 0.1 % bovine serum albumine (8076.2, Carl Roth, Germany) in PBS was used as control treatment for all recombinant proteins.

### Wnt reporter assay

HEK293T cells were seeded in a 96-well dish and were transfected with 20 ng TCF firefly luciferase (TOP-FLASH) plasmid and 2 ng CMV Renilla luciferase plasmid using ScreenFect® A according to the manufacturer’s protocol (ScreenFect GmbH, Germany). After 24 hours cells were treated with purified Wnt ligands and incubated overnight. Cells were harvested in 1x Passive Lysis Buffer (Promega, Germany) and processed according to the manufacturer’s protocol. TOPFLASH luciferase values were normalized to control Renilla luciferase values. Mean values (± SD) were calculated from four independent experiments.

### Western blotting

Secreted Wnt10b protein was detected by western blotting upon enrichment using Blue Sepharose beads (17-0948-01, Th. Geyer Inc., Germany). Enriched Wnt10b was resolved on 10 % SDS PAGE gels and detected by western blotting using anti-Wnt10b antibodies (MABN717, 1:1000, 5A7, Merck-Millipore, Germany, RRID: AB_3675944). Anti-HSC70 antibodies (#sc-7298, 1:1000, B-6, Santa Cruz, USA, RRID: AB_627761) were used as loading control.

### Cell cycle analysis

Cell cycle distribution and mitotic content was determined by FACS analysis using a BD FACSCanto™ II (BD Biosciences, Germany) and analyzed using FACSDiva™ software (Version 6, BD Biosciences), as described (Ertych et al, 2014).

### Measurements of microtubule plus-end growth rates

To measure microtubule plus-end assembly rates, comets of fluorescently labelled microtubule end-binding protein 3 (EB3) were tracked by live cell fluorescence microscopy. Cells were transfected with pEGFP-EB3 (kindly provided by Linda Wordeman, USA) or pcDNA3-EB3-StayGold (kindly provided by Atsushi Miyawaki, Japan) (Hirano M et al, 2022) plasmids and analyzed using a DeltaVision Elite live cell microscope as previously described (Bohly et al, 2022, Ertych et al, 2014). For mitotic measurements, cells were accumulated in mitosis by treatment with 2 µM dimethylenastrone (#SML0905, Sigma-Aldrich, Germany). Average assembly rates were calculated from 25 individual measurements per cell and a total of 30 cells from three independent experiments were analyzed per condition.

### Cell synchronization

To synchronize cells at the G1/S transition of the cell cycle, cells were synchronized by a double thymidine block protocol as described (Schmidt et al, 2021). 2 mM thymidine (#sc-296542A, Santa Cruz, USA) was added for 16 h. Cells were released into fresh culturing medium by washing with fresh culturing medium every 5 min for 30 min, cultured for 8 h and subjected to a second thymidine block for 16 h before releasing from the G1/S block. Cells were further analyzed at different time points after the release.

### Treatment of cell cycle synchronized cells

Cell cycle phase specific treatments (Fig. 2) were done using cells synchronized at the G1/S transition using a double thymidine protocol. Cells were released for various times as indicated in Fig. 2A and treated with DKK1, GSK3i, Wnt3a or Wnt10b for two hours followed by washout of the reagents. Treatments specific to early S phase (Fig. 3) were achieved by release from G1/S synchronization for two hours followed by washout of the treatments. Induction of replication stress in early S phase (Fig. 5 and 6) was achieved in the same manner while treatment with hydroxyurea was done two hours after the release from the G1/S block to avoid cell cycle arrest before cell enter S phase. Hydroxyurea was washed out four hours later.

### Detection of chromosome missegregation

The appearance of lagging chromosomes during anaphase was used as a measure for chromosome missegregation as described previously (Cimini et al, 2001). Cells grown on glass cover slips were enriched by cell cycle synchronization using a double thymidine block and release protocol coverslips, fixed with 2 % p-formaldehyde/PBS for followed by treatment with ice-cold methanol at -20°C for for 5 min. Mitotic spindles, kinetochores and chromosomes were detected by immunofluorescence microscopy using anti-α-tubulin antibodies (1:700, sc-23948, Santa Cruz, USA; RRID:AB_628410), anti-Cenp-C antibodies (1:1,000, PD030, MBL, USA; RRID:AB_10693556) and Hoechst33342 (#H3570, Thermo Fisher Scientific, Germany; RRID:AB_3675235). Images were captured using a Leica DMI6000B microscope (Leica, Germany) equipped with a Leica DFC360 FX camera and Leica LAS-AF software (Leica, Germany). A lagging chromosome was defined as Cenp-C-positive DNA clearly separated from the polar DNA masses in late anaphase cells.

### Detection of chromosomal breaks

Chromosomal breaks were detected on metaphase chromosome spreads. Cells were treated with 2 µM dimethylenasteron for 3 h to enrich mitotic cells in prometaphase. Cells were harvested, resuspended in 40 % (v/v) RPM1-1640 medium (PAN Biotech, Germany)/60% H_2_O and fixed in 75 % methanol/25 % glacial acetic acid. Cells were suspended in 100% glacial acetic acid and dropped onto pre-cooled microscopy slides. The slides were dried, and DNA was stained with Giemsa solution (AppliChem, Darmstadt, Germany). Bright-field microscopy was performed using a Leica DM IL LED microscope (Leica, Wetzlar, Germany) equipped with a ODC832 camera (Kern & Sohn GmbH, Balingen-Frommern, Germany) to detect and to quantify the appearance of chromosomes with visible breaks.

### Molecular DNA combing

Molecular DNA combing was performed to determine DNA replication fork progression rates and inter-origin distances as a measure for activation of additional origins (Moore et al, 2022). Newly synthesized DNA was sequentially labelled with 100 μM 5-Chloro-2ʹ-deoxyuridine (CldU; #C6891, Sigma-Aldrich, Germany) and 100 μM 5-Iodo-2ʹ-deoxyuridine (IdU; #I7125, Sigma-Aldrich, Germany) for 30 min each. Cells were harvested and processed using the FiberPrep DNA extraction kit (Genomic Vision, France) according to manufacturer’s protocol. The isolated and purified ssDNA was combed on salinized microscope slides (Genomic Vision, France) using a Molecular Combing System (Genomic Vision, France). Combed DNA samples were stained with primary anti-CldU antibodies (#ab6326, 1:10, BU1/75, ICR1, Abcam, UK, RRID:AB_305426), anti-IdU antibodies (#347580, 1:10, B44, BD Biosciences, Germany, RRID:AB_10015219), anti-ssDNA antibodies (#autoanti-ssDNA, 1:5, DSHB, USA, RRID:AB_10805144) and secondary antibodies conjugated to Alexa Fluor 488 (A21121, 1:25, Invitrogen, Germany, RRID:AB_2535764), Alexa Fluor 594 (#150160, 1:25, Abcam, UK, RRID:AB_2756445), and BV421 (#563846, 1:25, BD Biosciences, Germany, RRID:AB_2738449). Labelled DNA fibers were imaged using a Delta Vision Elite microscope (Delta Vision, SA) equipped with a PCO edge sCMOS camera (PCO, USA) at 63x magnification and labelled stretched of DNA were anlyzed to calculate fork speeds and inter-origin distances as described (Bohly et al, 2022).

### Statistical analysis

Statistical analysis was performed and graphs were drawn using GraphPad Prism 9.0 software (GraphPad Software, USA). Mean values and standard deviations (SD) were calculated for each experiment. Microtubule polymerization rate measurements and DNA combing analyses were depicted as box plots, where the whiskers represent the 10^th^ and 90^th^ percentile, the boxes the 25^th^ and 75^th^ percentile, the line in the boxes the median value and the plus sign within the boxes the average value. All experimental results are based on at least three independently performed biological replicates as indicated in the figure legends. Statistical analysis was performed using unpaired two-tailed *t*-tests. The statistical analysis of the Wnt/β-Catenin signaling activity was performed using and unpaired one-sample *t*-test using Microsoft Excel. Significances for experiments are indicated as p-values: ns (not significant): p ≥ 0.05, ∗: p < 0.05, ∗∗: p < 0.01, ∗∗∗: p < 0.001.

## Data availability

All raw data generated in this study will be made available. This study did not generate data that were deposited in public databases.

## Acknowledgments

We thank Atsushi Miyawaki, Feng Zhang and Linda Wordeman for providing plasmids. This work was funded by the Deutsche Forschungsgemeinschaft (DFG), project number 331351713 – SFB1324 (H. Bastians, G. Davidson, M. Boutros, S. Acebron). We acknowledge support by the Open Access Publication Funds of the Göttingen University.

## Author Contributions

Experiments were carried out by A.H. and F.W., J.W. performed Wnt reporter assays and *WNT10B* knockout cells were generated by O.V.. Experiments were analyzed and interpreted by A.H., F.W. J.W., G.D., O.V., M.B., S.A. and H.B. H.B. supervised the research and S.A. co-supervised parts of the research. The manuscript was written by H.B. and A.H. and all authors discussed results and commented on the manuscript.

## Conflict of interest

The authors declare no competing interests.

## Notes

### Competing Interest Statement

The authors have declared no competing interest.

### Summary of Updates

The order of the authors was incorrect and has been now corrected with Alexander Haass as first author and Holger Bastians as last and corresponding author.

