## Supplemental Figures and legends for "Wnt10b signaling regulates replication stress-induced chromosomal instability in human cancer"

#### Supplementary Figure legends

##### Figure S1 - HCT116 *WNT10b* knock-out cells

**(A)** Representative western blot detecting Wnt10b protein in cell culture supernatant from HCT116 and HCT116-*WNT10B* knock-out cells. **(B)** Wnt reporter assay in HEK293T cells after treatment with 400 ng/ml recombinant Wnt3a or Wnt10b protein for 24 hours. Data were normalized to control treatment for each replicate (mean ± SD, n=4, one-sample *t*-test).

##### Figure S2 – Cell cycle progression in HCT116 and HCT116-*EVI/WNTLESS* knockout cells

Representative cell cycle profiles of HCT116 and HCT116-*EVI*/WNTLESS knock-out cells after release from a G1/S cell cycle block. Cell cycle distribution was determined by FACS analysis**.**

##### Figure S3 – Wnt inhibition does not affect cell cycle progression

**(A)** Simultaneous determination of replication fork speed and inter-origin distances in cells upon treatment with DMSO or 100 nM of aphidicolin in the presence of absence of Wnt10b (RPE1-hTert) or upon Wnt inhibition using DKK1 treatment (HCT116). Box plots show mean values based on measurements of n>300 fibers per condition for fork speed and n>50 fibers per condition for inter-origin distances (two-tailed *t*-test).

**(B)** Experimental outline for the analysis of the cell cycle progression in HCT116 and HCT116-*EVI*/WNTLESS knock-out cells. **(C)** FACS-based quantification of the proportion of the indicated cells in different cell cycle stages after release from a G1/S cell cycle block (10,000 cells per condition, n=3).

### Figure S1

A

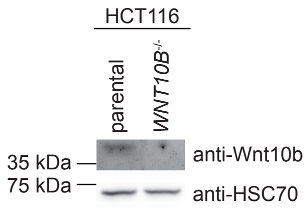
HCT116
parental
WNT10B-/-
35 kDa
75 kDa
anti-Wnt10b
anti-HSC70

B

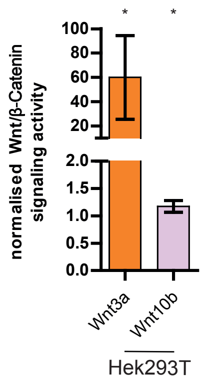
normalised Wnt/β-Catenin signaling activity
100
80
60
40
20
2.0
1.5
1.0
0.5
0.0
*
*
Wnt3a
Wnt10b
Hek293T

### Figure S2

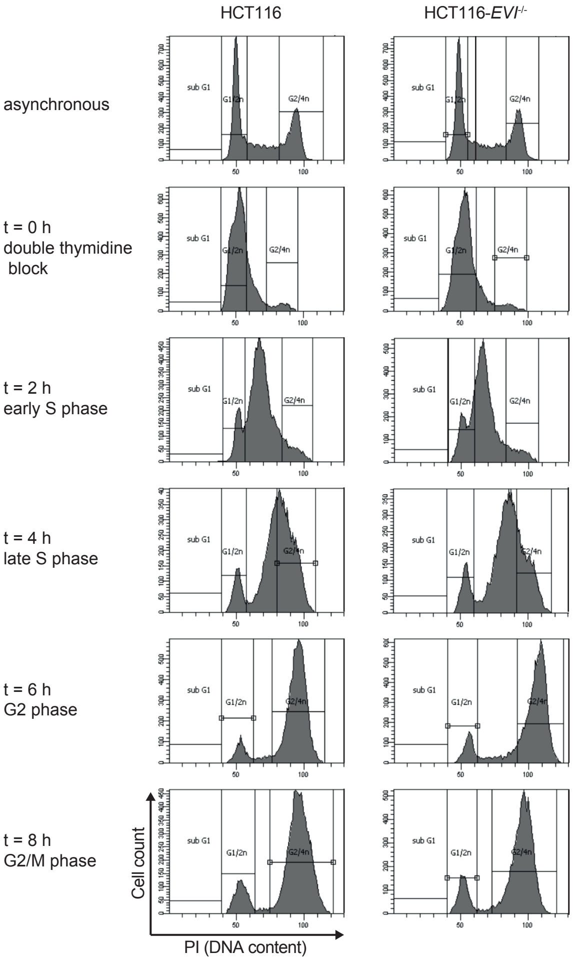
HCT116
HCT116-EVI-/-
asynchronous
t = 0 h
double thymidine
block
t = 2 h
early S phase
t = 4 h
late S phase
t = 6 h
G2 phase
t = 8 h
G2/M phase
sub G1
G1/2n
G2/4n
Cell count
PI (DNA content)

### Figure S3

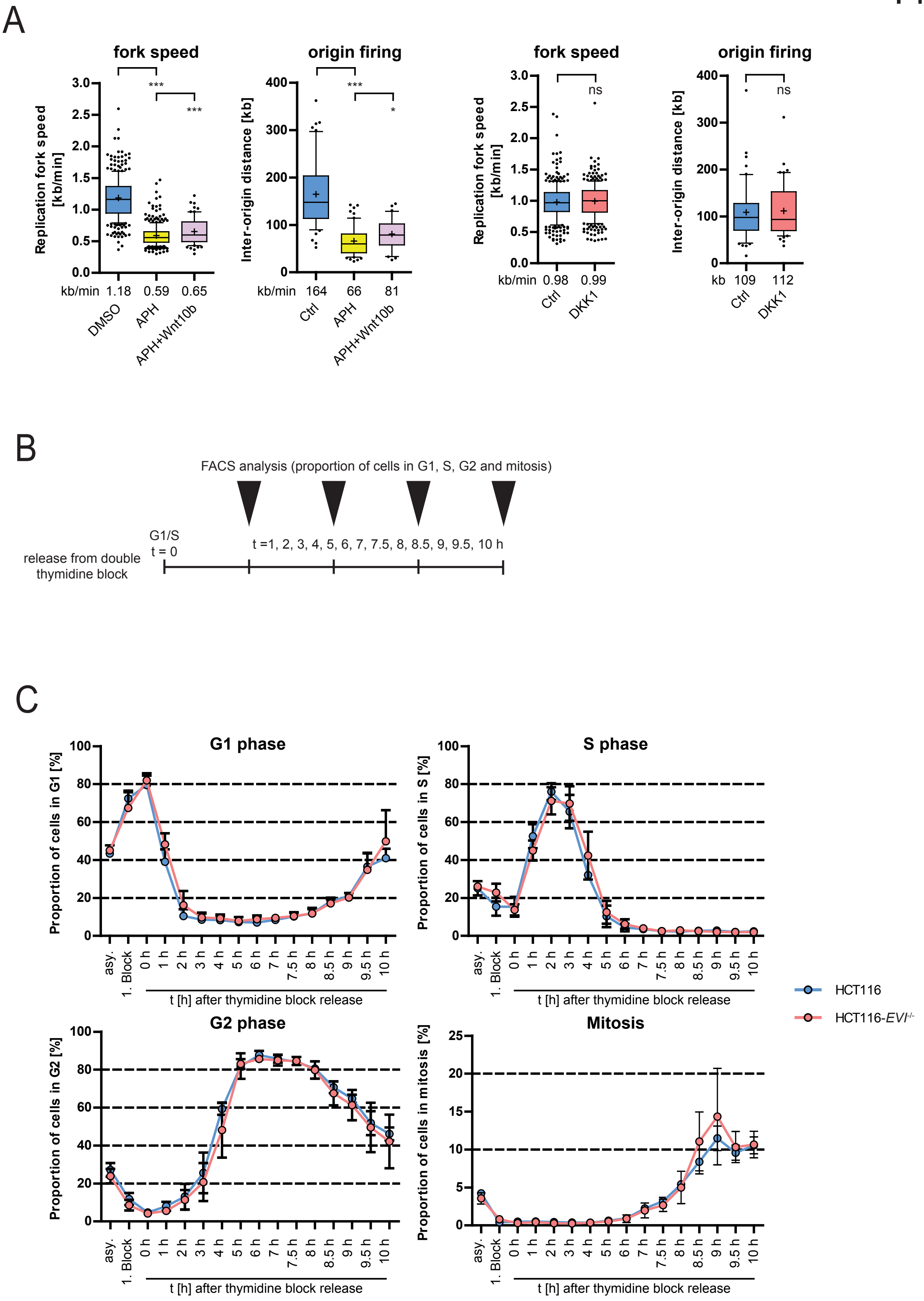
A
fork speed
origin firing
Replication fork speed [kb/min]
Inter-origin distance [kb]
***
*
ns
kb/min 1.18 0.59 0.65
DMSO APH APH+Wnt10b
kb/min 164 66 81
Ctrl APH APH+Wnt10b
kb/min 0.98 0.99
Ctrl DKK1
kb 109 112
Ctrl DKK1
B
FACS analysis (proportion of cells in G1, S, G2 and mitosis)
G1/S
t = 0
t =1, 2, 3, 4, 5, 6, 7, 7.5, 8, 8.5, 9, 9.5, 10 h
release from double thymidine block
C
G1 phase
S phase
G2 phase
Mitosis
Proportion of cells in G1 [%]
Proportion of cells in S [%]
Proportion of cells in G2 [%]
Proportion of cells in mitosis [%]
asy. 1. Block 0 h 1 h 2 h 3 h 4 h 5 h 6 h 7 h 7.5 h 8 h 8.5 h 9 h 9.5 h 10 h
t [h] after thymidine block release
HCT116
HCT116-EVI-/-
